# Helminthic parasite infections of endangered western hoolock gibbons (*Hoolock hoolock*) in degraded habitats of Bangladesh

**DOI:** 10.1101/2024.10.18.619014

**Authors:** M Tarik Kabir, Susan Lappan, Rajib Acharjee, Jesmin Sultana Pomy, Shahrul Anuar Mohd Sah, Nadine Ruppert

**Affiliations:** School of Biological Sciences, Universiti Sains Malaysia, 11800 Pulau Pinang, Malaysia; Malaysian Primatological Society, Taman Nuri, 08000 Kulim, Kedah, Malaysia; Department of Anthropology, Appalachian State University, Boone, NC, United States of America; Department of Zoology, University of Chittagong, Chattogram-4331, Chattogram, Bangladesh

**Keywords:** Nematodes, One Health, Parasitic Prevalence, Zoonosis, Small Apes

## Abstract

Although habitat degradation and anthropogenic disturbance can increase the risk of parasitic infections for many animals, including primates, little information about parasitism in wild gibbons is available, and almost nothing is known about the prevalence of helminth infection in wild populations. Globally threatened western hoolock gibbons (*Hoolock hoolock*, Harlan 1834) are exposed to significant anthropogenic disturbances across their range. We examined faecal samples from wild gibbons in three highly degraded and fragmented forests in Bangladesh: Lawachara National Park, Saltila forest, and Sheikh Jamal Inani National Park, for helminths, and identified two helminthic parasites, *Bunostomum* sp., and *Enterobius* sp. Overall helminthic parasite prevalence (percentage of samples containing parasites) and intensity (mean number of parasites individuals per sample) in faecal samples from wild gibbons were 82% and 12.1, respectively. This is the first report of *Bunostomum* sp. and *Enterobius* sp. infections in wild individuals of western hoolock gibbons or any other small ape. Habitat fragmentation, degradation, human encroachment, and other anthropogenic disturbances in Bangladesh may contribute to higher infection risk and transmission of zoonotic diseases in this highly arboreal gibbon species, with unknown consequences for their health and long-term survival. As parasitic infections can be transmitted within and across species, affecting humans, other primates, and other wild and domesticated animals, habitat protection and restoration and public awareness campaigns applying the One Health approach should be used to mitigate the spread of zoonotic disease in shared landscapes.

**HIGHLIGHTS:**

- First reported helminthic parasite infections with *Bunostomum* sp. and *Enterobius* sp. in wild gibbons
- Out of 50 collected fecal samples from the endangered Western hoolock gibbon in three degraded forests in Bangladesh, 41 (82%) tested positive for helminthic infections
- Parasite intensity ranged from 1 to 45 individuals per gram, with a median intensity of 9
- Lower parasite intensity (≤5) was more common in winter than in the monsoon season indicating seasonal variations

GRAPHICAL ABSTRACT

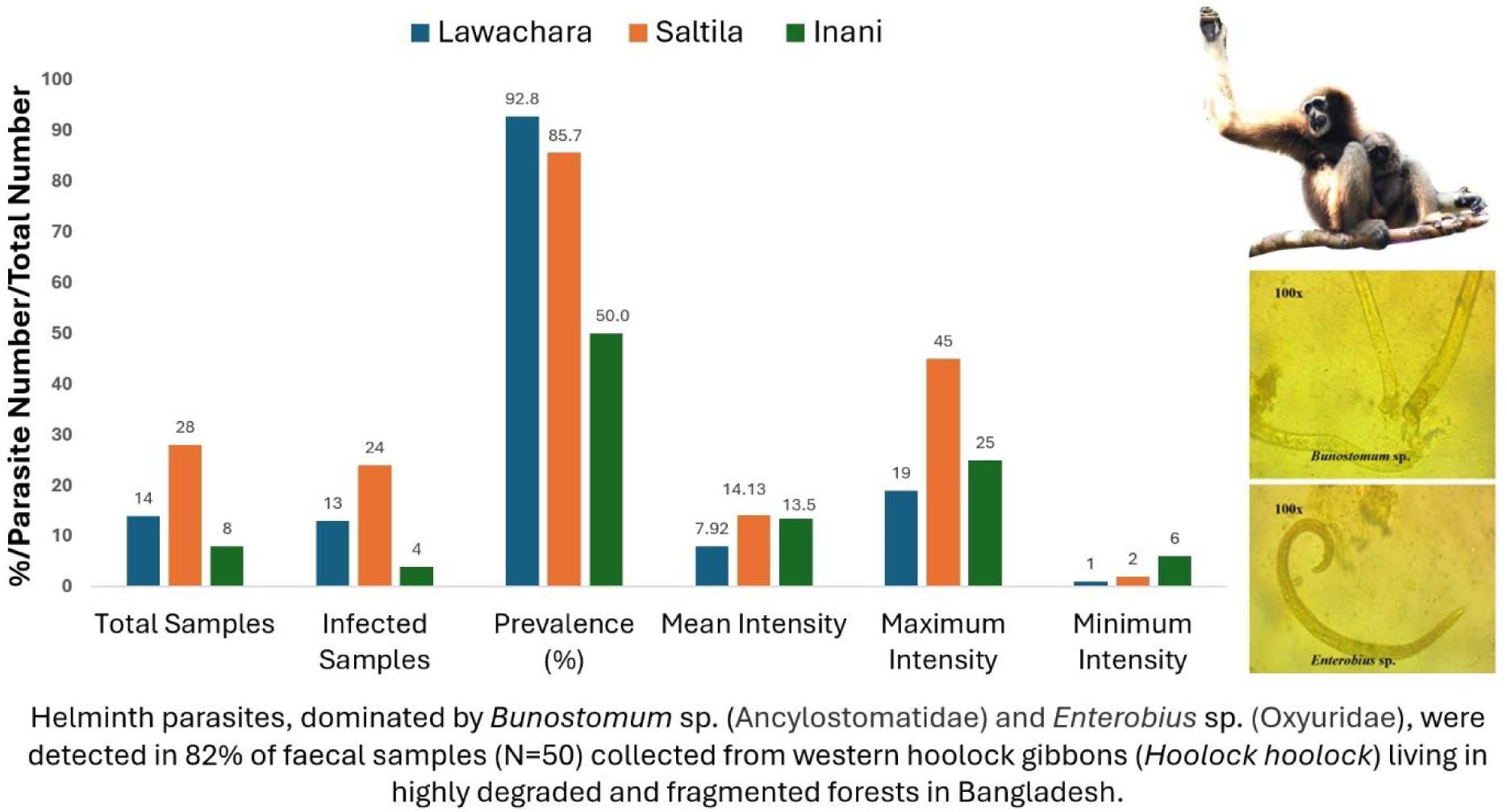

## 1. Introduction

Parasitism is ubiquitous in the animal kingdom, having independently evolved several times in at least 15 animal phyla (Weinstein and Kuris, 2016), and parasitic relationships can be important in structuring biodiversity (Poulin and Morand, 2000). Parasitic infections can affect the health, behaviour, ecology, demography, and evolution of individual species (Solórzano-García and Pérez-Ponce de León, 2018), and can shape the community structure of wildlife populations through their effects on trophic interactions, food webs, and intra– and interspecies competition (Preston and Johnson, 2010). Non-human primates (hereafter: primates) act as hosts of a comparatively wide spectrum of parasites (Kuntz, 1982), but little is known about the occurrence or prevalence of parasites in most wild populations.

Protozoa, helminths, and ectoparasites are major groups of parasites that can infect humans and primates. Most severe infections affecting primates are caused by helminths, including nematodes, cestodes, and trematodes (Toft and Eberhard, 1998). Captive primates are more likely to be infected with parasites transmitted through contact, including faecal-oral transmission, than free-living individuals because they are confined in smaller spaces, often in high densities (Herrera et al., 2019; Vonfeld et al., 2022), but wild animals may also become infected with parasites encountered in their environments (Herrera et al., 2019). Infection with parasites can result in severe health damage, and may also contribute to the risk of secondary infections due to the effects of infection on immune system functioning and nutrient uptake (Fekete and Kellems, 2007). Beyond the harm that parasitic infections can cause to the animals themselves, infected primates also have the potential to transmit parasites to humans in areas of ecological overlap due to our phylogenetically close relationship (Gómez et al., 2013). Thus, understanding the dynamics of parasite infection in anthropogenic landscapes is of particular importance both for the conservation of threatened animals and because of potential impacts on human welfare.

Higher parasite prevalence and species richness have been reported in primate populations sharing anthropogenic environments with humans. For example, lion-tailed macaques (*Macaca silenus*) in fragmented forests near human settlements in the Anamalai Hills, Western Ghats, India had higher parasite prevalence and were infected with a greater number of taxa than those living in fragments without adjacent human settlements (Hussain et al., 2013). Forest fragmentation was also linked to increased gastrointestinal parasite infection in western chimpanzees (*Pan troglodytes verus*) (Sá et al., 2013). Rhesus macaques (*M. mulatta*) in densely urban areas in Kathmandu, Nepal had a high prevalence of infection with intestinal parasites, and were infected by a diverse range of protozoans and helminths (Adhikari et al., 2023).

Habitat disturbance may also increase primate risk of infection with parasites due to changes in exposure to edge habitats in anthropogenic landscapes. Anthropogenic disturbance and habitat fragmentation can elevate the risk of parasitic infections and facilitate the transmission of zoonotic diseases among wildlife species and humans (White and Razgour, 2020). Habitat disturbance can also lead to stress and poor nutrition, which may act synergistically with elevated risk of exposure, leading to increased parasite loads (Borg et al., 2014;), and higher parasite diversity in infected individuals (Deb et al., 2014). For example, parasitic prevalence, species richness, and magnitude of multiple infections were greater among redtail guenons (*Cercopithecus ascanius*), red colobus (*Piliocolobus tephrosceles*) and black-and-white colobus (*Colobus guereza*) in the logged forest than undisturbed forest within Kibale National Park, Uganda (Gillespie et al., 2005), and groups of red colobus and black-and-white colobus living on habitat edges had a higher proportion of multiple infections and a higher prevalence of specific pathogens than groups living in the forest interior (Chapman et al., 2006).

The western hoolock gibbon (*Hoolock hoolock*; Hylobatidae, small apes) is the only gibbon species in Bangladesh and India, and its distribution range extends eastward into Myanmar (Brockelman et al., 2019). Due to severe habitat loss and fragmentation, the western hoolock gibbon is listed as Critically Endangered in Bangladesh (Feeroz et al., 2015) and Endangered globally (Brockelman et al., 2019). In Bangladesh, gibbons occur only in the mixed-evergreen hill forest fragments of northeast and southeast Bangladesh (Feeroz et al., 2015). The remaining population in Bangladesh is highly fragmented, and many populations are isolated in small, degraded forest fragments (Molur et al., 2005). Anthropogenic pressures on the remaining gibbon habitats in Bangladesh include agricultural activities, cattle grazing, illegal resource harvesting, betel leaf cultivation, lemon orchard, and local *jhoom* cultivation (a traditional shifting cultivation method using slash and burning) (Chowdhury et al., 2014; Reza and Hasan, 2019). Degradation of gibbon habitats may lead to food stress, resulting in poor body condition and reduced immune system functioning, making exposed animals overall more susceptible to different types of infections (Deb et al., 2014). The gibbon populations of Bangladesh are also likely to be at a high risk of parasitic infection as they share their habitats and forest resources with humans and livestock, which are potential reservoirs of pathogens in heavily anthropogenic landscapes.

Little is known about the prevalence of gastrointestinal parasites in wild gibbons. Ten species of parasites were detected in the faeces of wild white-handed gibbons (*Hylobates lar*) in Khao Yai National Park in Thailand, including two species of protozoa, one species of trematodes (*Dicrocoeliidae*) and seven species of nematodes (*Necator* sp., *Trichostrongylus* sp., *Strongyloides fuelleborni*, *Ternidens* sp., *Trichuris* sp., *Ascarius* sp., and an unknown species (Gillespie et al., 2013). The prevalence of parasites in this sample ranged from 4.4% to 91.3%, and the parasites detected included several that can cause substantial pathology and even death in primates (Gillespie et al., 2013). In a subsequent study of the same population, adult worms of the nematode *Streptopharagus pigmentatus* were also detected in white-handed gibbon faeces (Barelli and Huffman, 2017). Kharismawan et al. (2022) reported infection by unknown strongylids, *Trichostrongylus* sp., *Strongyloides* sp., and *Trichuris* sp. in wild Javan gibbons (*Hylobates moloch*) from Central Java, Indonesia, and Yalcindag et al. (2021) documented infection by *Oesophagostomum spp*. nematodes in wild white-bearded gibbons (*H. albibarbis*) in Indonesian Borneo. Studies in captive western hoolock gibbon populations have detected infections with several potentially pathogenic parasites, including *Balantidium coli*, *Trichuris* sp., hookworm eggs, whipworm eggs, *Giardia*, and oocysts of an unknown species (Muangkram et al., 2006; Nath et al., 2012; Raja, 2012), indicating their vulnerability to infection. However, captive animals are subjected to different exposure types and risks than wild animals, and no information is available on endoparasitic infections of wild western hoolock gibbons or other *Hoolock* species.

We assessed helminth parasite infections in faecal samples from wild western hoolock gibbons in three anthropogenically disturbed landscapes in northeast and southeast Bangladesh. Specific goals of this study were to identify helminth parasites infecting gibbons at these degraded sites, to compare the prevalence and intensity of parasite infection in gibbons across the three sites, and to assess the potential effects of season, age-sex class, severity of habitat fragmentation, and intensity of anthropogenic habitat disturbance on the prevalence and intensity of parasite infection in western hoolock gibbons in Bangladesh. This is the first attempt to identify helminth parasites or to assess the prevalence or intensity of parasite infection in wild western hoolock gibbons, and as such should provide valuable baseline data for health monitoring in these threatened populations.

## 2. Materials and methods

### 2.1. Study areas

Faecal samples were collected from the wild gibbons at Lawachara National Park (24.325440°N, 91.787036°E) and Saltila forest (24.169214°N, 91.407888°E) in northeast Bangladesh, and Sheikh Jamal Inani National Park (hereafter “Inani”, 21.228032°N, 92.068139°E) in southeast Bangladesh (Fig. 1). The study sites are characterized by a moist tropical climate with highest precipitation from April to September and a drier period from November to March (Uddin and Hassan, 2010).

**Figure 1:**
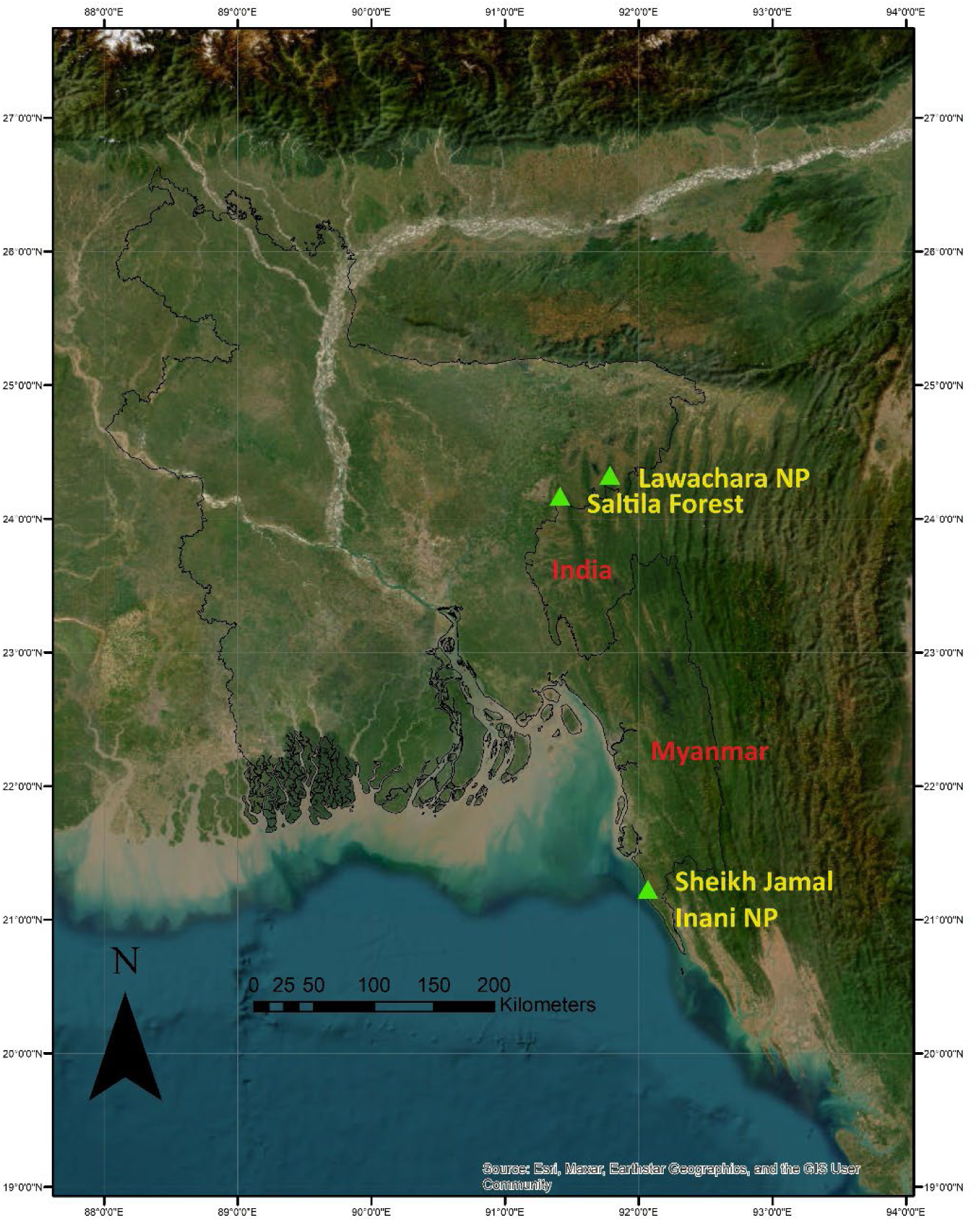
Locations of the study sites in Bangladesh. The black line indicates the borders of Bangladesh with Myanmar and India.

Lawachara is an isolated forest covering an area of 12.5 km^2^ and supporting an estimated population of 40 individuals (Naher et al., 2021). The group density at Lawachara was ca. 1.69 group/km^2^ (Naher et al., 2021). This area includes degraded forest interspersed with patches of secondary natural forests and plantation forests (Uddin and Hassan, 2010). Major tree species here are *Aphanamixis polystachia, A. chaplasha*, *Castanopsis tribuloides, Chukrassia tabularis, Dillenia pentagyna*, *Dipterocarpus turbinatus*, *Elaeocarpus floribundus*, *Ficus racemosa, Lagerstroemia parviflora, Lophopetalum fimbriatum, Quercus spicata, Steriospermum personatum, Toona ciliata, Vitex peduncularis* and *Xylia dolabiformis* (Uddin and Hassan, 2010). Threats at Lawachara include illegal logging, forest fires during the dry season, firewood collection, betel leaf cultivation, such as roads and tourism facilities, and uncontrolled mass tourism (Uddin and Hassan, 2010).

Saltila is a 13.55 km^2^ fragmented forest that is part of the ca. 26 km^2^ Raghunandan Reserved Forest. Within Saltila, only a small patch of disturbed secondary forest (ca. 1 km^2^) supports a remnant population of three individual gibbons (Kabir et al., 2024). Vegetation of the Saltila forest is comprised of small areas of natural forest, mixed vegetation, plantation forest, bamboo patches, and grasslands (Hosen and Ahamed, 2017). *Artocarpus chama, A. lacucha, Chickrassia tabulari, Garcinia zanthopyga, Ficus benghalensis, F. nervosa, F. benjamina, Prunus ceylonica,* and *Syzygium cumin* are the major plant species in Saltila (Hosen and Ahamed, 2017). Firewood collection and cattle grazing are the major observed anthropogenic disturbances at Saltila (MT Kabir, pers. obs.).

Inani covers an area of almost 80 km^2^ and constitutes a highly degraded and disturbed gibbon habitat without upper canopy trees (Kabir et al., 2021). Here, Kabir et al. (2021) reported the presence of seven gibbon groups (18 individuals). Major plant species at Inani include *Albizia* sp., *Amoora wallichii*, *Aphanamixis polystachya*, *Artocarpus chama*, *Bombax* sp., *Dillenia pentagyna*, *Duabhanga grandiflora*, *Ficus* sp., *Gmelia arborea*, *Hopea odorata*, *Lagerstroemia* sp., *Lannea coromandelica*, *Mesua ferrea*, *Magnifera longipes, Swintonia floribunda*, *Syzygium* sp., and *Tetrameles nudiflora*. Encroachment, illegal logging, expanding *jhoom* cultivation, and betel leaf vineyards are the major observed threats in Inani (Kabir, 2017).

### 2.2. Faecal sample collection

A total of 50 faecal samples were collected from six gibbon groups (Lawachara: three groups; Saltila: one group; Inani: two groups) during the monsoon and winter seasons from August 2022 to December 2023 (Table 1). Monsoon samples were collected from April to November and winter samples from December to March. Only freshly defecated samples were collected to avoid contamination. One group of gibbons in Lawachara and the Saltila group were sampled in both the moonson and winter seasons, while the other two Lawachara groups and the Inani groups were sampled only in a single season each.

**Table 1:** Description of the gibbon (*Hoolock hoolock*) study groups and sampling, including age-sex categories, group size, and number of faecal samples collected each season.

| Group number | Group composition |  |  |  |  | Total individuals | Seasons of sample collection |  | Total samples |
| --- | --- | --- | --- | --- | --- | --- | --- | --- | --- |
|  | Adult male | Adult female | Sub-adult | Juvenile | Infant |  | Monsoon | Winter |  |
| Lawachara 1 | 1 | 1 | 1 | - | - | 3 | 3 | 3 | 6 |
| Lawachara 2 | 1 | 1 | 2 | 1 | 1* | 6 |  | 5 | 5 |
| Lawachara 3 | 1 | 1 | 1 | - | - | 3 | 3 |  | 3 |
| Saltila 1 | 1 | 1 |  | - | - | 2 | 18 | 10 | 28 |
| Inani 1 | 1 | 1 |  | - | - | 2 | 4 |  | 4 |
| Inani 2 | 1 | 1 | 1 | - | - | 3 | 4 |  | 4 |
| **Total** |  |  |  |  |  |  | **32** | **18** | **50** |

\*sample was not collected from the infant

The ability to continuously follow the gibbons affected the frequency of faecal sample collection. The gibbons at Saltila were habituated, which allowed the research team to collect more samples and to sample more consistently at Saltila than at Lawachara and Inani. Continuous following of the gibbons at Inani was almost impossible due to the difficulty of the terrain and the heavy use of the landscape by Asian elephants.

When samples were collected, we noted the weather, time, date, and age-sex class of the gibbon that produced each sample (Klaus et al., 2018). Gibbon individuals were categorized as infant, juvenile, sub-adult, adult male, or adult female based on coat color, sexual morphology, and behaviour patterns following Ahsan (1994) and Kakati et al. (2009). Anthropogenic disturbances were categorized as agricultural activites (*i.e.,* planting crop fields, lemon orchards, and betel leaf vineyards), road infrastructure, human settlements, and cattle grazing. Anthropogenic disturbances were visually observed and recorded with GPS waypoints. After mapping the locations of these disturbances, distances from these locations to the home ranges of the gibbon groups were calculated in Google Earth (Appendix 2).

### 2.3. Helminth assessment

Freshly defecated faecal samples were carefully removed from the ground with a clean spoon to avoid contamination and kept in a sterilized 50 ml falcon tube. Each falcon tube was labeled with the group identity, time of collection, site of collection, date of collection, and sample number. About 15-20 grams of faeces were preserved in 70% ethanol and used for analysis (Gillespie et al., 2006). To avoid further deterioration of the samples, the samples were preserved in the refrigerator for a maximum of two weeks at 4°C before analysis. Samples were prepared for microscopic analysis by faecal floatation and sedimentation (Gillespie et al., 2006).

Prepared microscopy slides were studied under both binocular (XSZ-107BN) and trinocular microscopes (Euromex Oxion Inverso) at various magnifications (100x and 400x) to resolve morphological characters for identification of helminths to the genus or, where possible, species level. For each identified taxon, the parasitic prevalence and intensity were measured in each sample (see Shaw et al., 2018). Parasitic prevalence is the proportion of host individuals infected with a particular parasite. We calculated prevalence as the percentage of gibbons infected with a particular parasite or eggs (Bush et al., 1997). Intensity is the average number of individual parasites per gram of each sample (Shaw et al., 2018).

### 2.4. Data analysis

Because the relationship between distance to roads, settlements, and agricultural fields and human activity in the forest is not well understood, and our sample sizes did not permit a detailed exploration of these relationships, we used group identity, rather than specific habitat variables, as an indicator of the underlying habitat quality in each group’s home range for data analysis.

We used model selection with generalized linear mixed models (GLMM) to estimate the effects of season, adult sex, and group identity on parasite prevalence (Model 1) and parasite intensity in infected adults (Model 2). Because sampling was unbalanced, we used the Satterthwaite approximation with robust estimation. For parasite prevalence, which involves a binary outcome variable (infected/not infected), we used a binomial distribution with a logit link. Parasite intensity is a count variable, so we used a Poisson probability distribution and a log link in the GLMM for intensity. Individual identity was included as a random factor in both models because most individuals were sampled repeatedly. Because sample sizes for juvenile and subadult individuals were not adequate to determine whether age class affected parasite prevalence or intensity, we included only adult samples in the models. For each analysis, we used the Akaike Information Criterion corrected for small sample sizes (AICc) to compare candidate models with each other and with a null model that had the same error structure and included no predictor variables. We used ΔAICc (the difference between the AICc for a candidate model and the AICc for the model with the lowest AICc) to determine which were the best models, selecting the candidate models with ΔAICc≤2.

## 3. Results

Overall, 50 faecal samples were collected and assessed for infection with helminthic parasites, and 41 samples (82%) were positive for helminthic infections (Table 1). Disturbance was high at all sites, with all gibbon groups living ≤1.1 km from the nearest agricultural field, ≤3.5 km from the nearest road, and ≤2.5 km from the nearest settlement. All sites showed evidence of grazing activity (Table 2).

**Table 2:** Distance of each study group from the nearest location showing each type of anthropogenic disturbance.

| Location | Gibbon group | Mean intensity of parasites in each group | Distance of nearest anthropogenic disturbance (km) |  |  |  |
| --- | --- | --- | --- | --- | --- | --- |
|  |  |  | Agricultural field | Roads | Human settlements | Grazing |
| Lawachara | Group 1 | 8 | 0.55 | 0.11 | 0.85 | 0 |
| Lawachara | Group 2 | 3.8 | 0.6 | 0.3 | 0.77 | 0 |
| Lawachara | Group 3 | 14 | 0.3 | 0 | 0.9 | 0 |
| Saltilla | Group 5 | 12.1 | 2 | 3.5 | 2.5 | 0 |
| Inani | Group 6 | 13.5 | 0 | 1.3 | 0 | 0 |
| Inani | Group 7 | 0 | 1.1 | 2.8 | 1.2 | 0 |

The parasite fauna in the samples was dominated by *Bunostomum* sp. (Ancylostomatidae) and *Enterobius* sp. (Oxyuridae) (Fig. 2). Another unidentified parasite was found in one sample from Inani (Fig. 2). Infections with *Bunostomum* sp. (n = 41 samples), *Enterobius* sp. parasites (n = 2) or *Enterobius* sp. eggs (n =1) were identified at all three study sites, but parasite prevalence and intensity of *Bunostomum* sp. were different across sites (Fig. 3). Parasite prevalence was highest at Lawachara (92.8%), followed by Saltila (85.7%) and Inani (50.0%) (Fig. 3).

**Figure 2:**
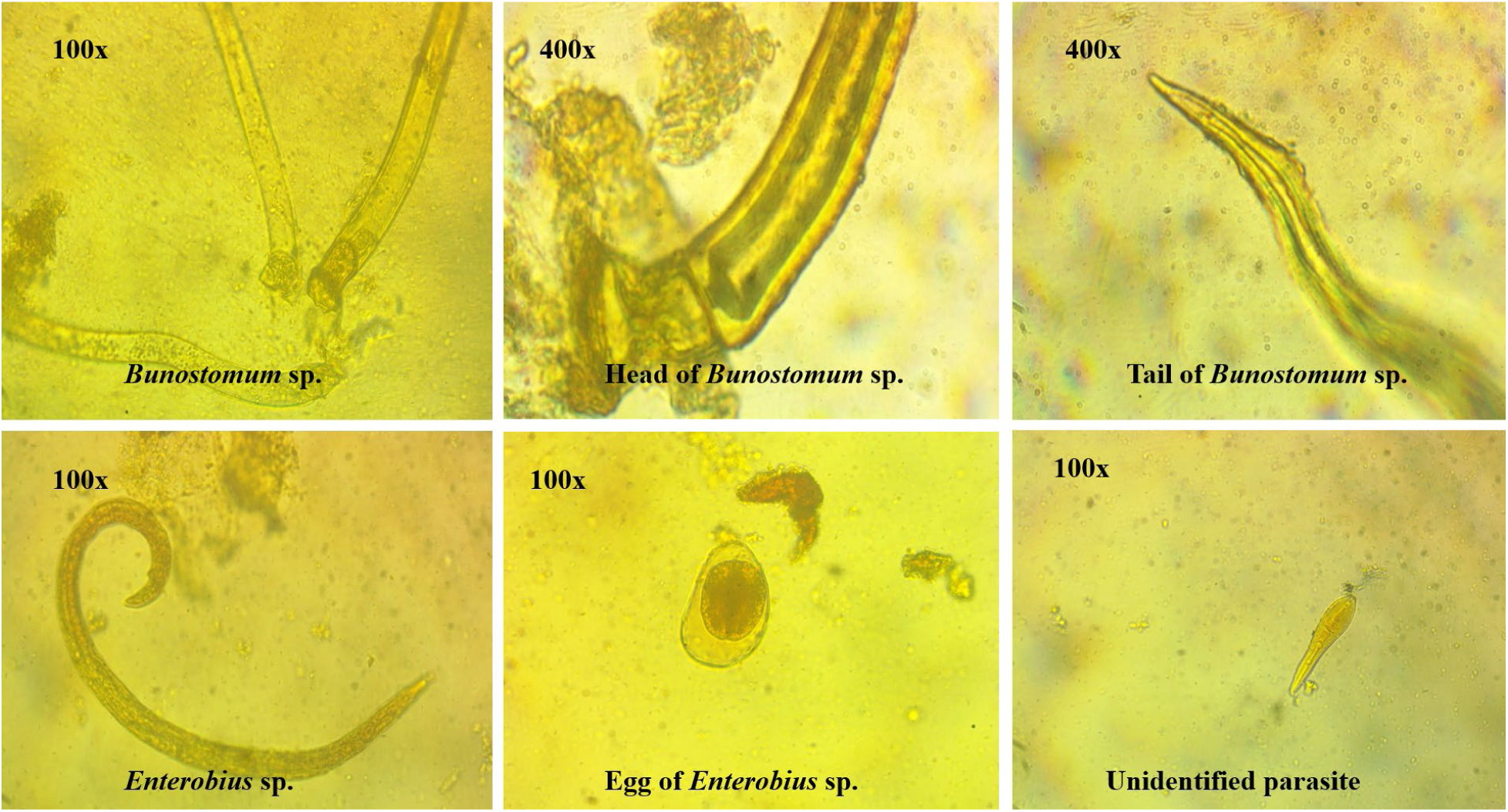
Parasites detected in western hoolock gibbons’ (*Hoolock hoolock*) faecal samples.

**Figure 3:**
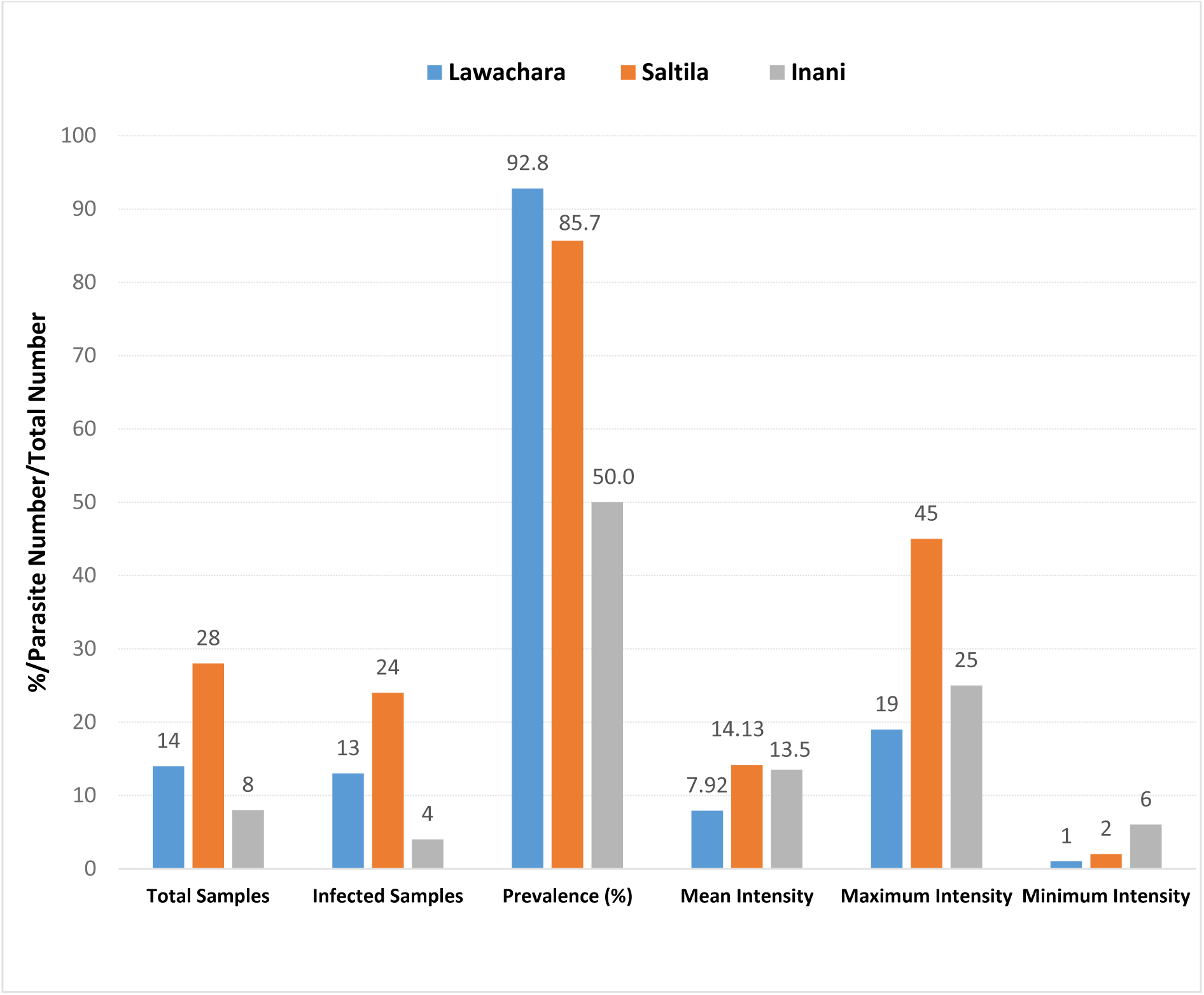
Epidemiological parameters of infected gibbons (*Hoolock hoolock*) groups at the three study sites.

Most individuals and most groups produced at least one sample infected with parasites. However, parasites were not detected in any individual in the ‘Inani 2’ group or in the juvenile in ‘Lawachara 2’, which was the only juvenile sampled in this study. In Saltila, parasites were found in most, but not all, samples in both seasons (Table 3). No single-variable model for parasite prevalence in adults performed better than the null model (Table 3). Models with multiple explanatory variables could not be built due to the low variance in parasite prevalence. Parasite intensity ranged from one to 45 individuals per gram, with a median intensity of nine. In infected samples, values ≤5 for parasite intensity were more common in winter (6 of 16 samples, 37.5%) than in the monsoon (2 of 24 samples, 8.3%) and were detected in only three groups: ‘Lawachara 1’ (2 of 6 samples, 8.3%), ‘Lawachara 2’ (2 of 4 samples, 50%), and ‘Saltila’ (4 of 24 samples, 16.7%) (Table 3). Few adult samples had parasite intensity ≤5 (adult females: 1 of 21 samples; adult males: 2 of 15 samples), but most infected samples from subadults had intensity ≤5 (3 of 5 samples). The two models that best-predicted parasite intensity (ΔAICc≤2) in adults both included season and group and one also included adult sex (Table 4). No other model had ΔAICc≤2 (Table 5). The top four models all included season, and the confidence intervals around the estimates for season in all models did not include zero, confirming the importance of season as a predictor of parasite intensity. Despite the selection of models including group and sex, the estimates for these variables included zero in all selected models. Parasite prevalence was the same for the male and female adults in each of the groups in Inani and Lawachara, but the female gibbon in Saltila had a slightly higher parasitic prevalence than the male gibbon in both seasons (Table 3).

**Table 3.** Parasite prevalence and intensity (mean±SD) in six western hoolock gibbon (*Hoolock hoolock*) groups in degraded habitats in Bangladesh. Individuals with parasite prevalence ≤50% or parasite intensity ≤5 in a season are bold.

| Group | Season | Sex | Parasite prevalence (N) | Parasite intensity (N) |
| --- | --- | --- | --- | --- |
| Inani 1 | Monsoon | Female | 100±0% (3) | 16.0±7.8 (3) |
|  |  | Male | 100% (1) | 6 (1) |
|  | Winter | Female | - | - |
|  |  | Male | - | - |
| Inani 2 | Monsoon | Female | **0% (1)** | - |
|  |  | Male | **0% (1)** | - |
|  |  | Subadult | **0% (1)** | - |
|  | Winter | Female | - | - |
|  |  | Male | - | - |
|  |  | Subadult | - | - |
| Lawachara 1 | Monsoon | Female | 100% (1) | 8 (1) |
|  |  | Male | 100% (1) | 9 (1) |
|  |  | Subadult | 100% (1) | **1 (1)** |
|  | Winter | Female | 100% (1) | 8 (1) |
|  |  | Male | 100% (1) | 11 (1) |
|  |  | Subadult | 100% (1) | **5 (1)** |
| Lawachara 2 | Monsoon | Female | - | - |
|  |  | Male | - | - |
|  |  | Subadult | - | - |
|  |  | Juvenile | - | - |
|  | Winter | Female | 100% (1) | 6 (1) |
|  |  | Male | 100% (1) | **4 (1)** |
|  |  | Subadult | 100±0% (2) | **4.5±2.1 (2)** |
|  |  | Juvenile | **0% (1)** | - |
| Lawachara 3 | Monsoon | Female | 100% (1) | 7 (1) |
|  |  | Male | 100% (1) | 16 (1) |
|  |  | Subadult | 100% | 19 (1) |
|  | Winter | Female | - | - |
|  |  | Male | - | - |
|  |  | Subadult | - | - |
| Saltila | Monsoon | Female | 91.7±11.3% (12) | 16.4±11.3 (11) |
|  |  | Male | 66.7±51.6% (6) | 17.8±15.0 (4) |
|  | Winter | Female | 100±0% (3) | 18.3±11.4 (3) |
|  |  | Male | 85.7±38% (7) | 5.5±5.1 (6) |

**Table 4.** Results of the GLMM model (binary logistic) selection procedure for parasite prevalence in hoolock gibbon faecal samples with season, group, and sex as factors. Individual identity was included as a random factor. Reference categories were male sex, winter season, and Lawachara group 2. Estimates were calculated using the Satterthwaite approximation and robust estimation and are shown with 95% confidence intervals.

| Model | AICc | ΔAICc | Factor | β | SE | Lower | Upper |
| --- | --- | --- | --- | --- | --- | --- | --- |
| Null | 208.905 | 0 |  |  |  |  |  |
| Sex | 210.905 | 2 | Female | 1.054 | 1.2607 | -1.492 | 3.600 |
| Season | 211.620 | 2.715 | Monsoon | 1.430 | 1.1991 | -0.992 | 3.852 |
| Group | 318.245 | 109.340 | Inani 1 | 0.0000 | 2078.07 | -4210.56 | 4210.56 |
|  |  |  | Inani 2 | -31.132 | 2190.48 | -4469.46 | 4407.19 |
|  |  |  | Lawachara 1 | 0.0000 | 2078.07 | -4210.56 | 4210.56 |
|  |  |  | Lawachara 3 | 0.0000 | 2399.54 | -4861.94 | 4861.94 |
|  |  |  | Saltila | -13.771 | 1696.73 | -3451.68 | 3424.14 |

**Table 5.** Results of the GLMM model (Poisson, log link) selection procedure for parasite intensity in hoolock gibbon faecal samples with season, sex, and group as factors. Individual identity was included as a random factor. Reference categories were male sex, winter season, and Inani group1. Estimates were calculated using the Satterthwaite approximation and robust estimation and are shown with 95% confidence intervals.

| Model predictors | AICc | ΔAICc | Factor | $\beta$ | SE | Lower | Upper |
| --- | --- | --- | --- | --- | --- | --- | --- |
| Season and group | 195.205 | 0 | Monsoon | 0.346 | 0.1219 | 0.097 | 0.595 |
|  |  |  | Lawachara 1 | -0.084 | 0.4049 | -1.197 | 1.029 |
|  |  |  | Lawachara 2 | -0.483 | 0.4986 | -1.604 | 0.637 |
|  |  |  | Lawachara 3 | -0.017 | 0.4221 | -1.116 | 1.081 |
|  |  |  | Saltila | 0.278 | 0.3723 | -0.922 | 1.478 |
| Season, sex, and group | 195.971 | 0.766 | Monsoon | 0.342 | .1221 | 0.092 | 0.592 |
|  |  |  | Female | 0.142 | 0.2884 | -0.754 | 1.037 |
|  |  |  | Lawachara 1 | -0.048 | 0.4474 | -1.430 | 1.335 |
|  |  |  | Lawachara 2 | -0.456 | 0.5333 | -1.745 | 0.834 |
|  |  |  | Lawachara 3 | 0.020 | 0.4642 | -1.315 | 1.355 |
|  |  |  | Saltila | 0.311 | 0.4178 | -1.229 | 1.851 |
| Season | 200.411 | 5.206 | Monsoon | 0.384 | 0.1173 | 0.147 | 0.623 |
| Season and sex | 201.179 | 5.974 | Monsoon | 0.378 | 0.1178 | 0.138 | 0.617 |
|  |  |  | Female | 0.139 | 0.2464 | -0.461 | 0.740 |
| Group | 209.382 | 14.177 | Lawachara 1 | -0.228 | 0.4313 | -1.379 | 0.924 |
|  |  |  | Lawachara 2 | -0.815 | 0.5083 | -1.974 | 0.344 |
|  |  |  | Lawachara 3 | -0.007 | 0.4506 | -1.149 | 1.135 |
|  |  |  | Saltila | 0.164 | 0.4019 | -1.042 | 1.371 |
| Sex and group | 210.136 | 14.931 | Female | 0.176 | 0.3005 | -0.724 | 1.075 |
|  |  |  | Lawachara 1 | -0.193 | 0.4640 | -1.595 | 1.210 |
|  |  |  | Lawachara 2 | -0.788 | 0.5361 | -2.109 | 0.533 |
|  |  |  | Lawachara 3 | 0.030 | 0.4828 | -1.328 | 1.388 |
|  |  |  | Saltila | 0.195 | 0.4365 | -1.331 | 1.772 |
| Null | 216.603 | 21.398 |  |  |  |  |  |
| Sex | 217.189 | 21.984 | Female | 0.173 | 0.2910 | -0.419 | 0.764 |

## 4. Discussion

To the best of our knowledge, our results detected the first confirmed observation of helminthic parasite infection of western hoolock gibbons in the wild, and the first confirmation of infection with *Bunostomum* sp. or *Enterobius* sp. in any wild small ape (Hylobatidae; Table 6).

**Table 6.** Parasites detected in previous studies of wild gibbons (Hylobatidae), with host species, location, and reported prevalence.

| Type | Family | Parasite species | Gibbon species | Prevalence | Location | References |
| --- | --- | --- | --- | --- | --- | --- |
| Nematoda | Ancylostomatidae | *Bunostomum* sp. | *Hoolock hoolock* | 82% | Bangladesh | This study |
|  |  | *Necator* sp. | *Hylobates lar* | 8.7% | Thailand | Gillespie et al., 2013 |
|  | Ascarididae | *Ascaris* sp. | *Hylobates lar* | 8.7% | Thailand | Gillespie et al., 2013 |
|  | Onchocercidae | *Brugia malayi* | *Hylobates lar* | - | Malaysia | Laing et al., 1960 |
|  |  | *Brugia pahangi* | *Hylobates lar* | - | Malaysia | Laing et al., 1960 |
|  | Oxyuridae | *Enterobius* sp. | *Hoolock hoolock* | 6% | Bangladesh | This study |
|  | Spirocercidae | *Streptopharagus pigmentatus* | *Hylobates lar* |  | Thailand | Barelli & Huffman, 2017 |
|  | Strongylidae | *Ternidens* sp. | *Hylobates lar* | 91.3% | Thailand | Gillespie et al., 2013 |
|  |  | *Oesophagostomum* sp. | *Hylobates albibarbis* | - | Indonesia | Yalcindag et al., 2021 |
|  |  | *Ternidens* sp./<br>*Oesophagostomum* sp. | *Hylobates moloch* | 25% | Indonesia | Kharismawan et al., 2022 |
|  |  | Unknown larval strongylid | *Hylobates lar* | 21.7% | Thailand | Gillespie et al., 2013 |
|  |  | Unknown strongylid | *Hylobates moloch* | 25% | Indonesia | Kharismawan et al., 2022 |
|  | Strongyloididae | *Strongyloides fuelleborni* | *Hylobates lar* | 56.5% | Thailand | Gillespie et al., 2013 |
|  |  | *Strongyloides* sp. | *Hylobates moloch* | 5.5% | Indonesia | Kharismawan et al., 2022 |
|  | Trichostrongylidae | *Trichostrongylus* sp. | *Hylobates lar* | 13.0% | Thailand | Gillespie et al., 2013 |
|  |  | *Trichostrongylus* sp. | *Hylobates moloch* | 25% | Indonesia | Karismawan et al., 2022 |
|  | Trichuridae | *Trichuris* sp. | *Hylobates lar* | 91.3% | Thailand | Gillespie et al., 2013 |
|  |  | *Trichuris* sp. | *Hylobates moloch* | 8.3% | Indonesia | Kharismawan et al., 2022 |
| Trematoda | *Dicrocoelidae* | *Dicrocoelidae* sp. | *Hylobates lar* | 8.7% | Thailand | Gillespie et al., 2013 |
| Protozoa | Balantiidae | *Balantidium coli* | *Hylobates lar* | 87% | Thailand | Gillespie et al., 2013 |
|  | Cryptosporidiidae | *Cryptosporidium* sp. | *Hylobates lar* | 4.4% | Thailand | Gillespie et al., 2013 |

*Bunostomum* sp. (82%) had a much higher prevalence in our sample than *Enterobius* sp. (6%). *Bunostomum* is a hookworm (Ancylostomatoidae), a superfamily of nematodes that feed on blood and parasitize the gastrointestinal tracts of many mammals (Seguel and Gottdenker, 2017).

While *Bunostomum* infections are most often reported in domesticated sheep, goats, buffaloes, and cattle, they also infect several families of wild ungulates (Ukashatu et al., 2012; Seguel and Gottdenker, 2017; Gunaithilaka et al., 2018; Sanda et al., 2019), and have been reported in wildlife in other mammalian orders, including primates (Seguel and Gottdenker, 2017).

*Bunostomum* infection has previously been recorded in several other South Asian primates, including wild lion-tailed macaques (*Macaca silenus*; Hussain et al., 2013), bonnet macaques (*M. radiata*, Kumar et al., 2018), rhesus macaques (*M. mulatta*; Adhikari et al., 2022), and Nilgiri langurs (*Trachypithecus johnii*; Tiwari et al., 2017). However, the prevalence of *Bunostomum* in western hoolock gibbons in this study was substantially higher than the reported prevalence in most other primate populations (wild lion-tailed macaques: 59%, Hussain et al., 2013; bonnet macaques: 1.5%, Kumar et al., 2018; Nilgiri langurs: 0.45%, Tiwari et al. 2017), but similar to *Bunostomum* prevalence in urban rhesus macaques in Nepal (87.6%; Adhikari et al., 2022. This may be due to the higher risk of exposure to these parasites in heavily degraded anthropogenic landscapes.

Hookworm infections generally occur through cutaneous exposure to or ingestion of infected soil. In Bangladesh, *Bunostomum* occurs in domesticated sheep, goats, and buffaloes (Khatun et al., 2021; Islam et al., 2022). Cattle grazing was observed at all three study sites during the study period, suggesting that the gibbons may have been exposed to *Bunostomum* through contact with soil infected by the faeces of grazing ruminants.

*Enterobius*, commonly known as pinworm, which we detected in 6% of samples, is the most frequently reported helminth genus infecting humans and primates (e.g., Atelidae: Stoner and Gonzalez Di Pierro, 2006; Cercopithecidae: Hugot, 1999; Hominidae: Hugot 1993; and Indridae: Muelenbein et al., 2003). Humans are a reservoir for the infection of *Enterobius* and bi-directional transmission between humans and primates has been documented (Strait et al., 2012). Previous studies of wild gibbons have detected infections with a variety of helminths, including *Ascarias* sp., *Brugia malayi, B. pahangi, Necator* sp., *Streptopharagus pigmentatus, Strongyloides* sp., *Ternidens* sp., *Trichostrongylus* sp., and *Trichuris* sp. (Liang et al., 1960; Gillespie et al., 2013; Barelli and Huffman, 2017; Kharishmayan et al., 2022; Table 6). Patterns of prevalence and species richness varied among taxa and studies. *Ternidens* and *Trichuris* had the highest prevalence (91.3%) in wild white-handed gibbons (*Hylobates lar*), followed by *Strongyloides fuelleborni* (56.5%), *Trichostrongylus* sp. (13.0%), *Necator* sp. (8.7%), *Ascaris* sp. (8.7%), besides unknown larval strongylides (21.7%; Gillespie et al., 2013). Nematode prevalence was lower (31.9%) among Javan gibbons (*H. moloch*) in Central Java, Indonesia (Kharismawan et al., 2022). Because of their strict arboreality, gibbons are often assumed to have lower exposure to soil-transmitted helminths than primates that routinely travel on the ground, but our results add to a growing body of evidence that suggests that the prevalence of some helminths in gibbons are comparable to or higher than those reported from other, more terrestrial, primate species (Solórzano-García and Pérez-Ponce de León, 2018; Medkour et al., 2020), however, species richness in intestinal parasites has been low across studies of wild gibbons relative to species richness in great apes (Gillespie et al., 2013). Although gibbons rarely descend to the ground, habitat destruction and fragmentation are driving animals to adapt and develop new feeding strategies, which can lead to an increased risk of parasitic infections (Deb et al., 2014). During the data collection, we observed gibbons in the highly degraded habitat at Inani coming down to the ground to feed on soil (Appendix 1; MT Kabir, pers. obs.), a behaviour often associated with mineral deficiencies (Pebsworth et al., 2018). Reduced availability of critical minerals for frugivores may compromise immune system functioning (Rode et al., 2006), making exposed individuals more susceptible to infection, which may amplify the effect of greater risk of parasite exposure.

This study showed that parasitic intensity was higher during the monsoon season. Seasonal variation in infection risk may occur for many reasons, including seasonal differences in parasite survival and development, seasonal changes in host behaviour that affect exposure risk, or seasonal changes in host physiology (Altizer et al., 2006; Shearer and Ezenwa, 2020). For nematodes, high humidity and temperatures are associated with faster development and reproduction (Turner and Getz, 2010). Parasitic prevalence in white-handed gibbons at Khao Yai National Park in Thailand was also higher during monsoon season (Gillespie et al., 2013; Barelli and Heistermann, 2013) and the prevalence of parasites in Tibetan macaques (*M. thibetana*) was higher during summer than winter (Yang et al., 2022). Our results, therefore, align with a growing body of data suggesting seasonal variation in risk of nematode infection in nonhuman primates.

For some animal populations, sex is another important predictor of parasite prevalence or intensity (Klein, 2004), a pattern that has reported in several nonhuman primate species (e.g., *Pongo abelii*: Mul et al., 2007; *Procolobus rufomitratus* and *Cercocebus galeritus*: Mbora and Munene, 2006; *Pan troglodytes verus*; Metzger, 2015). Sex differences in parasitism may occur because of sex-related behavioural differences that can affect exposure risk or due to physiological variation in the ability to mount an effective immunological response following exposure (Klein, 2004; Mariencheck, 2024). We found that parasite prevalence and intensity were usually similar for pair-mates, and our models did not provide strong support for an effect of sex on parasite prevalence or intensity, although sex was included in some selected models.

Since gibbon groups are small, and paired male and female gibbons generally travel together, forage in the same food resources, and share a home range, their risk of exposure to parasites may be very similar. There were slight differences in parasite prevalence between the male and female in the group at Saltila, with the female having a slightly higher prevalence of parasites in both seasons and a higher intensity in winter. This may reflect greater power to detect small differences for this group, which was sampled more intensively than the other groups, or it may reflect an unusual pattern of spatial interactions among the male and female in this isolated group. Although briefly meeting up for mating attempts, they ranged separately for most of the period of sample collection and had overlapping but distinct home ranges (Kabir et al., 2024).

The very high prevalence of nematode infection in our study suggests that heavy anthropogenic disturbance may increase the risk of nematode exposure or infection in wild primates. Sharing of wildlife habitats by people and livestock can increase the risk of infection with gastrointestinal parasites (Obanda et al., 2019), and habitat degradation is known to influence infection risk and risk of zoonosis (Tiwari et al., 2017; Wilkinson et al., 2018). Isolation in small fragments may affect local host population density, which is related to parasite prevalence and species richness for primates (Mbora and McPeek, 2009), and habitat disturbance may lead to changes in primate behaviour in ways that affect the risk of exposure. In landscapes where food availability has changed as a result of human activities, changes to primate nutrition and health may also affect the ability of the host immune system to respond effectively to parasite exposures (Rode et al., 2006; Deb et al., 2014). In this study, habitat fragmentation, anthropogenic disturbances, and sharing of resources between humans, livestock, and gibbons likely created opportunities for the transmission of parasites and promoted high prevalence and intensity at all three study sites. While we were not able to systematically evaluate the relationships between disturbance factors and parasite prevalence in our study groups, it is noteworthy that the home range of the infected gibbons at Inani was situated immediately adjacent to a human settlement area but the home range of the uninfected group was more than 1km away from the infected group and from the closest human settlement. However, both the prevalence and intensity of parasite infection in the group at Saltila were high, despite Saltila being the most distant of all of the study groups from the nearest settlement or agricultural field, which suggests that multiple factors are likely to affect parasite infection in western hoolock gibbons.

Gibbons have been reported to use antihelminthic and antimicrobial plant species for self-medication when infected with parasites (Barelli and Huffman, 2017). At our study site, *Alstonia scholaris* (family Apocynaceae) is a powerful medicinal native tree species as its leaves and flowers contain chemicals with antihelminthic and antimicrobial properties (Thankamani et al., 2011). During this study, western hoolock gibbons and Phayre’s langurs (*Trachypithecus phayrei*) were seen consuming leaves and flowers of *A. scholaris* in Saltila forest (M.T. Kabir, pers. obs.). While our data do not allow a more detailed examination of the relationship between parasitism and the use of potentially medicinal plants by wild primates, these observations highlight the importance of behavioural strategies, such as self-medication, in the management of disease risk in wild primates (Laumer et al., 2024), and the complexity of interspecies interactions in threatened habitats in Bangladesh.

Infection with gastrointestinal parasites can have severe health impacts. *Bunostomum* sp. can cause severe anaemia, reduced growth, tissue damage, severe inflammation, weight loss, and even death in infected animals (Barabosa et al., 2017; Seguel Gottdenker, 2017), so the detection of *Bunostomum* in threatened wildlife populations is concerning. Another soil-borne helminth, *Strongyloides* sp., that has been reported in wild white-handed gibbons (Gillespie et al., 2013; Kharismawan et al., 2022), can cause high mortality in captive gibbons (DePaoli and Johnsen, 1978). The detection of widespread nematode infections in the Critically Endangered western hoolock gibbon is therefore concerning, especially as it may compound the impacts of other threats to their persistence in the wild in Bangladesh and globally.

To mitigate this threat, spatial overlap in land use by gibbons, humans, and ruminants in shared habitats should be monitored and, where possible, grazing of ruminants in critical gibbon habitats should be avoided. Changes in human behaviour may also be necessary to reduce human-to-gibbon zoonosis. For example, Lawachara National Park is a popular tourist destination in Bangladesh, and heavy visitor pressure may increase the risk of transfer of human diseases to the gibbon population there. Therefore, clear guidelines for visitors to reduce the risk of disease transmission should be prepared and communicated to park staff, local tourism guides, and park visitors to reduce the impacts of tourism on wildlife. Captive primates generally have higher rates of infection than wild primates (Herrera et al., 2019; Vonfeld et al., 2022), so animals rescued from the illegal wildlife trade may also be potential sources of infection if they are released into wild populations without appropriate screening. A wildlife rescue center near Lawachara is currently rehabilitating two young gibbons for eventual reintroduction (https://www.thedailystar.net/news/bangladesh/news/hope-hoolock-gibbons-3208961), making further exploration of the parasitology of wild gibbons particularly urgent. To address the effects of poor habitat quality on parasite prevalence in anthropogenic landscapes, restoration, and enrichment of degraded gibbon habitats in Bangladesh with natural food plants may reduce behaviours such as consumption of soil, and may enhance gibbon nutrition and health. This may help to minimize immune system stress responses that can make gibbons more susceptible to parasite infections (Deb et al., 2014).

Ultimately, to address the risks that parasites can pose for these threatened gibbons and other sympatric mammals, including humans, the local Forest Department should include consideration of parasitic infections in their management strategies, by applying a holistic One Health approach for habitat protection, wildlife conservation, and tourism management in the future (Destoumieux-Garzon et al., 2018). Above all, existing anthropogenic disturbances inside the forests should be controlled and curbed to better protect Bangladesh’s last gibbons.

## Acknowledgments

We are grateful for the financial support from the Bangladesh Forest Department through their SUFAL Innovation Grant and from the Wildlife Conservation Network. We thank the Bangladesh Forest Department for permitting us to carry out the fieldwork in the eastern hill forests of Bangladesh. We are thankful to the Parasitology Laboratory of the University of Chittagong, Chattogram, Bangladesh for allowing the use of their facilities. Thank you to Mr. Razib Chandra Bhowmick, Department of Parasitology, Universitiy of Chittagong, who extended his support for the planning of this study during the initial stage of the project. We also acknowledge Mr. Ishtiaq Uddin Ahmad and Dr. M Farid Ahsan for their continuous encouragement to carry out this work. This work would not have been possible without the generous support from the numerous local field assistants, we owe you.

---

**Appendix 1:**
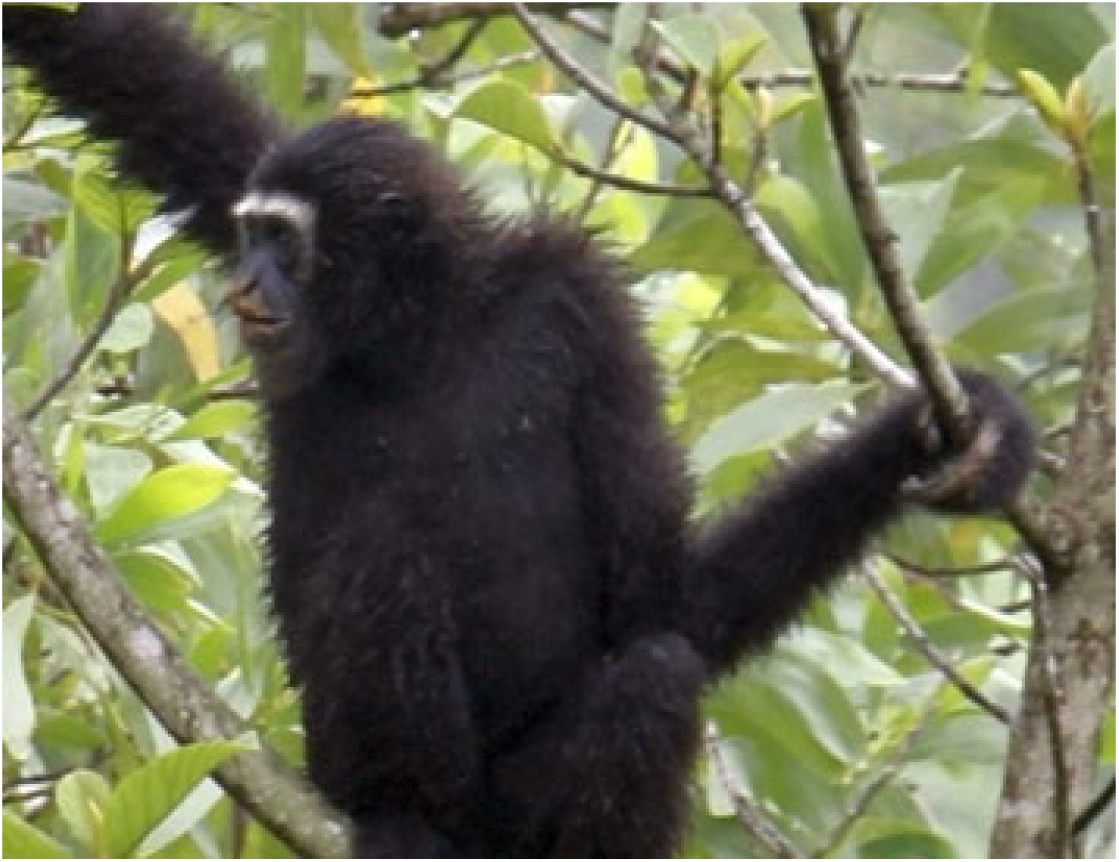
Infected male gibbon (*Hoolock hoolock*) with mud on his mouth and hand indicating consumption of soil at Inani, Bangladesh.

**Appendix 2:**
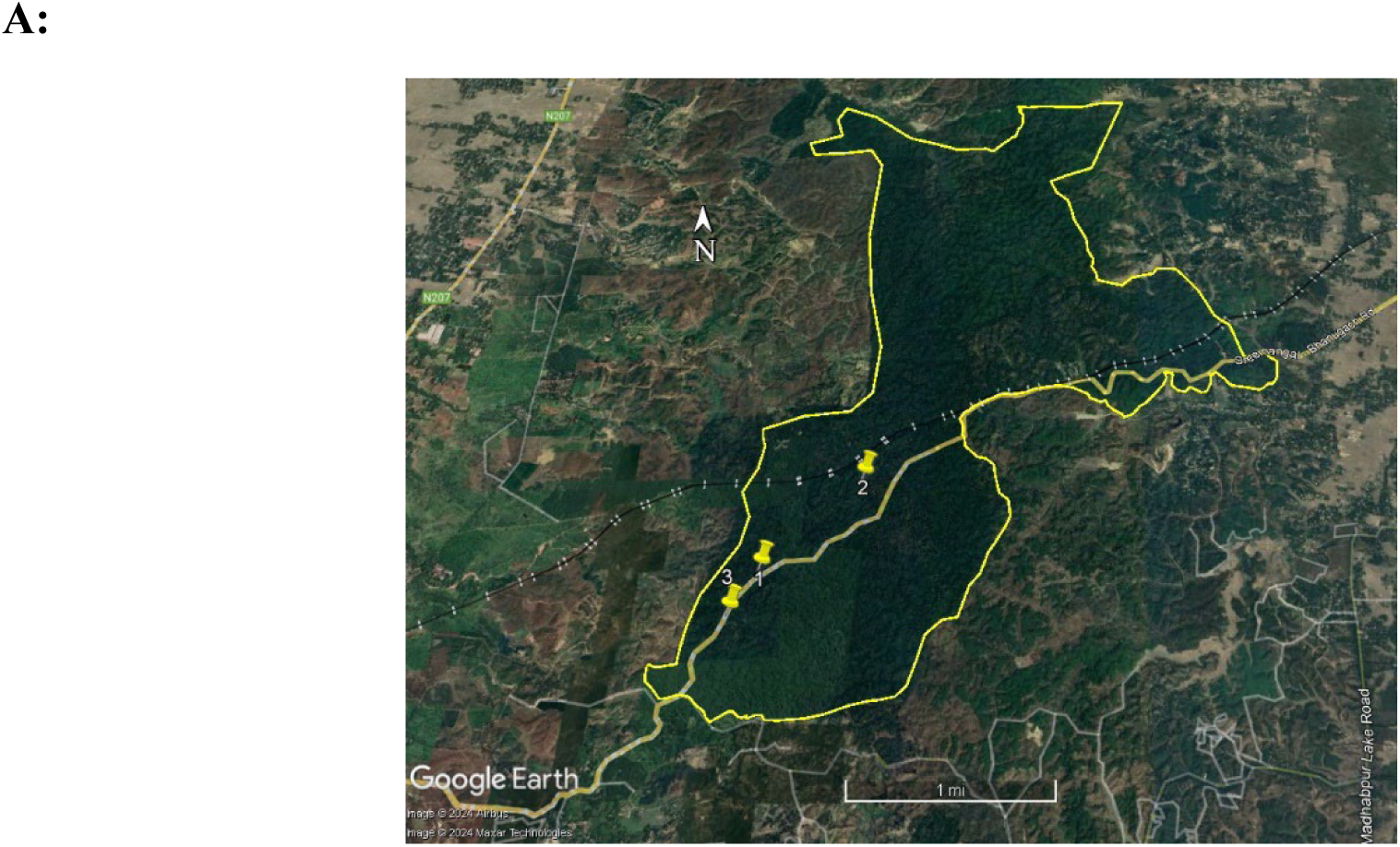
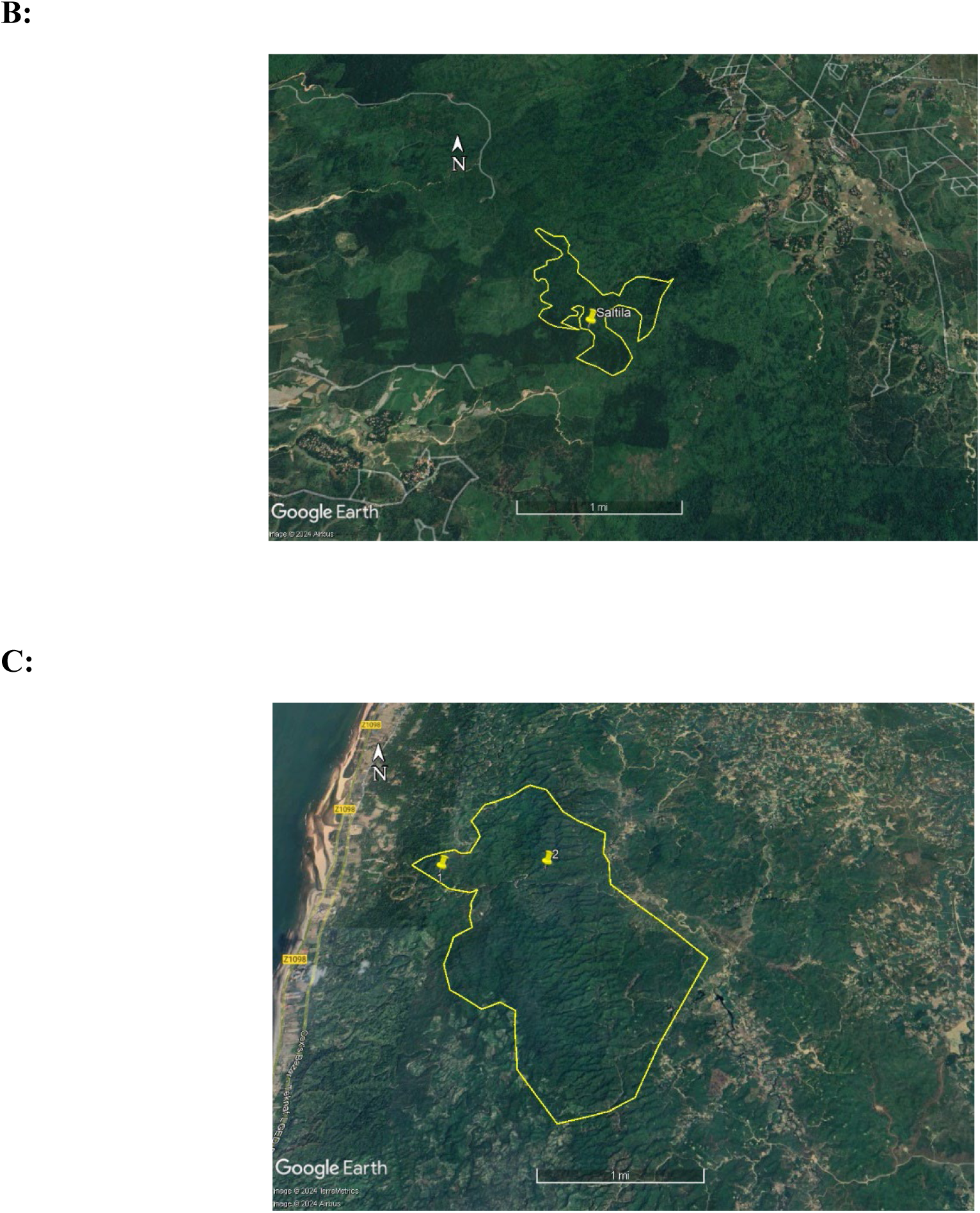
Locations of the sampled gibbon groups. A: Lawachara National Park. B: Saltila forest. C: Sheikh Jamal Inani National Park. Yellow placemarks show the location of the sampled gibbon groups. (Source: Google Earth)

